# Drug-bacterial membrane interactions: A Tale of two force fields

**DOI:** 10.1101/2023.09.27.559676

**Authors:** Subhadip Basu, Sandip Mandal, Prabal K. Maiti

## Abstract

Tuberculosis (TB), caused by *Mycobacterium tuberculosis* (M.tb), a gram-positive bacteria, is known to infect and kill millions of people worldwide, specifically in poor and developing countries. M.tb has unique characteristics in terms of the presence of an extraordinarily thick cell wall structure, mainly composed of mycolic acid (MA) and arabinogalactan (AG). Between these components, mycolic acid is of particular interest because it exhibits extremely low permeability for most of the drug molecules, thus helping M.tb to survive against medical treatment. Hence, it is extremely useful to study the interactions between drug molecules and mycolic acids in order to better design potential TB medicine. On that premise, the present work aims to computationally model mycolic acid monolayer and probe its permeability against some first-line TB drugs, namely ethambutol, ethionamide, and isoniazid. It is a known fact that such modelling strongly depends on the interaction parameters used. The current study addresses this issue by employing two widely used force fields-GROMOS 54A7-ATB (hereafter GROMOS) united atom force fields and CHARMM36 (hereafter CHARMM) all-atom force fields to collate the drug-mycolic acid interplay. Our findings indicate that both force fields provide consistent results in terms of the mode of drug-monolayer interactions but significantly differ for the monolayer permeability. Mycolic acid monolayer generally exhibited a higher free energy barrier of crossing using CHARMM FF, while GROMOS parameters provide better stability of drug molecules on the monolayer surface, which can be attributed to the greater electrostatic potential at the monolayer-water interface. Both the force field parameters predicted the highest resistance against ethambutol, the most soluble molecule. However, significant differences can be recorded for the other two molecules with the change in the interaction parameter set. In summary, our study provides insight into the dependency of *in silico* modelling of membranes on force field parameters together with the guidelines to investigate the drug-membrane interactions consisting of non-standard lipids/fatty acids.

## 1. Introduction

Tuberculosis (TB), the world’s top infectious killer, is responsible for ∼ 10.6 million infections and around 1.5 million deaths yearwise.^1^ Majority of TB infections and related deaths occurred in the poor and developing countries like India, China, Bangladesh, Nigeria, South Africa.^2^ Among these countries, India holds the largest share of the global TB infections.^1^ In 2019, more than 2.4 million TB cases were found in India.^3^ Although TB mainly infects the lungs, infections can be found in other parts, as well. TB is contagious and spread through air, when a patient with lung TB sneezes, coughs, or spits. Co-infections like HIV, emergence of multi-drug resistance TB, lack of healthcare infrastructure, and discontinuation of the treatment in the midway are some of the root causes behind the drastic global impact of TB.^2^

*Mycobacterium tuberculosis* (M.tb) is the causative agent of the infectious disease tuberculosis, and it belongs to the gram positive bacteria family, Mycobacteriaceae.^4^ It is distinct from most other gram-positive bacteria because of the presence of a extra thick cell wall structure located outside the peptidoglycan layer, found in gram-positive bacteria.^5^ This extraordinary thick cell wall structure contains long chain fatty acids, namely mycolic acids (MA), and polysaccharides called arabinogalactan (AG).^6^ Among these components, mycolic acids bestow M.tb with unique properties that defy medical treatment by lowering the efficacy of antibiotics/biocides, making the organism more resistant to chemical damage and dehydration.^7^ Mycolic acids are also responsible for the bacterium growth inside macrophages, effectively hiding it from the host immune system. Mycobacterial mycolic acids possess some distinct characteristics. They are longer, and contain 70-90 total carbon atoms, among which typically 24 carbon atoms belong to a fully saturated R-branch and the rest are part of the unsaturated mero chain.^8^ Usually two positions are there in the mero chain, that may be occupied by functional groups.^8^ The proximal position (nearer the α-hydroxy acid) contains exclusively cis– or trans-olefin or cyclopropane.^9^ However, the distal position may be the same as the proximal position or contain one of a variety of oxygen moieties such as R-methyl ketone, R-methyl methyl ether, methyl-branched ester, or R-methyl epoxide.^8^ M.tb cell wall contains three kinds of mycolates: α-mycolates, methoxymycolates, and ketomycolates, with relative abundance of 51%, 36% and 13%, respectively.^8^ The exact packing of the mycolic acids in the bacterium cell wall remains elusive. Previously, it was reported that, the mycolic acids can fold into three distinct “W”, “U” and “Z” conformers,^10^ although additional conformations have also been listed in the literature.^11^

Because of the important role of the mycolic acid in the drug resistance of M.Tb, it is of utmost importance to study the interactions of existing standard TB drugs or any potential TB drug candidate with the mycolic acid monolayers. Previously, Hong et. al. explored drug-mycolic acid interactions by computational means, they did not consider the anomalous diffusion of the drugs within the mycolic acid clusters, which usually takes place inside a tightly packed bio membrane.^9^ Against this background, we tried to compute the effective resistance and permeability of the mycolic acid monolayer against some of the well-known first line TB drugs, namely ethambutol, ethionamide, and isoniazid. For simplicity, we have modelled only α-mycolates, the abundant type found in M.Tb bacterial cell wall. One important aspect of modelling drug membrane interactions, specifically for a less explored fatty acid like mycolic acid, is the choice of the appropriate force field. In the present study, we have considered two commonly used force field families,CHARMM36 all-atom (CHARMM) and GROMOS 54A7-ATB (GROMOS), to model the drug-mycolic acid interactions to investigate the consistency of the results obtained from the both set of force fields, and also to probe the dependency of such modelling on the choice of interaction parameters.

## 2. Simulation Details

### 2.1 Modelling of mycolic acid-drug systems

α-mycolic acid was modelled using both all-atom and united atom representations. Packing of 100 α-mycolic acid molecules in a rectangular single monolayer was performed using MEMGEN and PACKMOL 20.3.1.^12,13^ 100 water molecules per mycolic acid chain were added to the simulation box, above and below the monolayer. For all-atom representations, the systems were modelled using CHARMM36 all-atom force field parameters (abbreviated as CHARMM) with CHARMM-modified TIP3P water model.^14–16^ The parameters were generated using CHARMM-GUI and CGENFF web interface.^17–21^ For the united atom representations, GROMOS 54A7-ATB (abbreviated as GROMOS) force field parameters, generated using Automated Topology Builder, with SPC/E water model was used to compute the molecular interactions.^22–24^ Initial configurations of the systems together with the structures of the molecules used for the study have been depicted in Fig. 1.

**Fig. 1:**
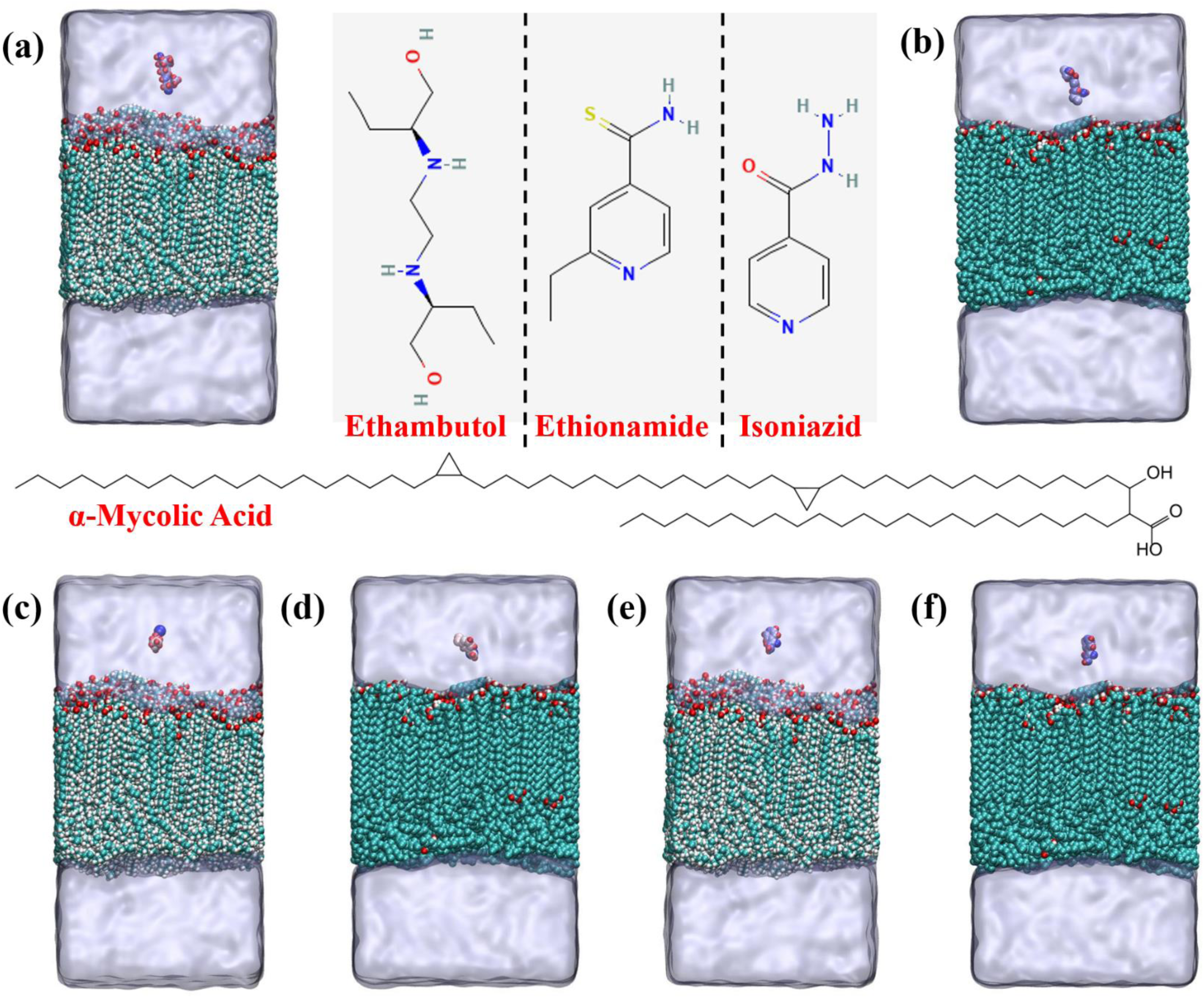
Molecular structures of drug molecules and initial configurations of the drug-monolayer systems. Structures of Tuberculosis drug molecules used in this study is shown in the upper-middle panel of the figure. The structure of α-mycolic acid is depicted in the middle. Initial configurations of ethambutol-monolayer system is presented for (a) CHARMM36 and (b) GROMOS 54A7-ATB FF. Starting configurations of ethionamide-monolayer system is shown in (c) and (d) for CHARMM36 FF, and GROMOS 54A7 FF, respectively. Snapshots of Isoniazid-monolayer systems at 0 ns is shown for (e) CHARMM36 and (f) GROMOS 54A7 FF. Colour code for mycolic acid: C:Pale Blue, O: Red, H: white. Colour code for Drug molecule is based on atomic mass. For ethambutol/isoniazid: H:Red, O: Blue, N: light blue, C:whitish blue. For ethionamide: S: Deep Blue, H: red, C: reddish white, N: white. Water is presented using ice-blue transparent surface.

The systems was subjected to energy minimization in 200,000 steps using steepest descent algorithm to remove any close contacts between atoms. The minimized system was subsequently subjected to a 10ns equilibrium run in NVT ensemble at 300K, followed by a 100ns run in NPT ensemble with 1 bar of pressure. The drug molecule (Ethambutol/Ethionamide/Isoniazid) was then added to the equilibrated system, 1.0 nm above the surface of the monolayer, replacing necessary number of water molecules. The drug molecules were placed on the side of the monolayer containing the –COOH groups of the α-mycolic acids (head region). The drug-monolayer systems were again energy minimized using steepest descent algorithm to remove any close contacts. The minimized systems were then again equilibrated in NVT ensemble for 1ns (at 300K), followed by a second phase of equilibration in NPT ensemble (at 1 bar pressure) for 10ns. Position restraints were put on the drug molecules during the equilibration phases. The equilibrated systems were then subjected to production run of 250ns and steered molecular dynamics simulation for either 1ns (for all-atom representation) or 750 ps (for united atom representation). Modified Berendsen thermostat with a coupling constant of 0.1 ps was used for temperature coupling, while the pressure was maintained using Parrinello-Rahman barostat with a coupling constant of 0.5 ps.^25,26^ Semi-isotropic coupling scheme was employed to account for the anisotropic structure of the mycolic acid molecules. Throughout the simulation time, all bonds involving hydrogen atoms were restrained using LINCS algorithm.^27^ A simulation box of 6.32×6.32×12.51 nm^3^ was used for the systems with CHARMM FF, while the dimension of the simulation box was 6.58×6.51×11.95 nm^3^ for GROMOS FF. Periodic boundary conditions were employed in all three directions with a cut-off distance of 1.2 nm to compute the short-range LJ interactions and short-range part of the coulomb interactions. Particle mesh Ewald (PME) summation technique was used to calculate long-range coulombic interaction.^28^ The integration time step used was 2fs and the trajectories of the systems were saved at an interval of 10ps for subsequent analysis. All simulations were performed in GROMACS-2022.3 package and the visualization was done using VMD 1.9.3 software.^29,30^ All of the analysis were carried out using gmx modules and in-house python scripts. We have studied a total of six drug-monolayer systems.

### 2.2 Steered MD simulations and Umbrella Sampling

Steered molecular dynamics simulations were performed to pull the drug molecule through the monolayer with a constant velocity of 0.01 nm/ps. During pulling, a harmonic spring with a spring constant of 1000 kJ mol^−1^nm^−2^ was applied on the drug molecule along the pulling direction, while the monolayer was taken as static reference. The trajectory was recorded for every 0.1 ps. In case of Isoniazid using all-atom representation (CHARMM36 force field), a spring constant of 1500 kJ mol^−1^nm^−2^ was used to keep the pulling motion steady.

Windows/configurations for the umbrella sampling was generated from the steered MD simulations at an interval of 0.1 nm along z-direction.10ns run was performed on each window to obtain the probability distribution of the systems for every configurations. For Isoniazid and CHARMM force field parameters, uneven sampling scheme (windows with 0.05 nm spacing for up to 1.5 nm distance from COM of monolayer and after that windows with 0.1 nm spacing were taken) was opted to ensure the overlap between the probability distributions of two consecutive sampling windows. PMF profile was computed using weighted histogram analysis method (WHAM) embedded in GROMACS, from the probability distribution obtained for each window.^31^ We have used umbrella sampling technique to further compute the local resistance (R(z)), effective resistance (R_eff_), and effective permeability (P_eff_) of the membrane in the following manner. First, Position dependent diffusion coefficient (D(z)) was calculated for every umbrella sampling window using the following equation:^32,33^

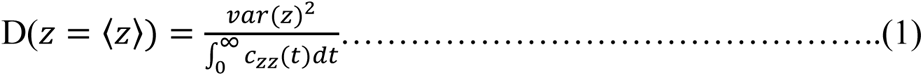

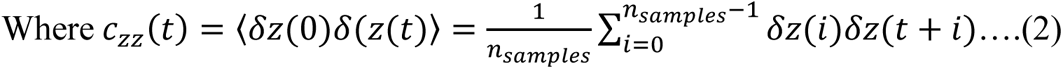

Local resistance value R(z) for each umbrella sampling window was obtained using equation (3), given below. ^32,33^

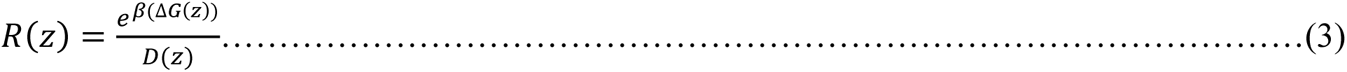

Here, ΔG or the free energy difference was obtained from the PMF curve at different values of z. Effective resistance and effective permeability of the monolayer was calculated by integrating eq. (3) and is given by eq. 4.

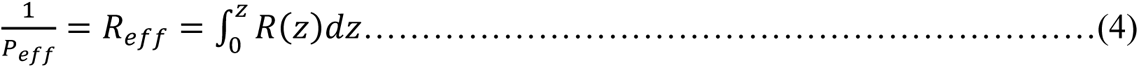

## 3. Results

In this section, the nature of the interactions between the mycolic acid membrane and the drug molecules will be illustrated using both CHARMM36 all atom FF and GROMOS 54A7-ATB FFs together with a description of the thermodynamic free energy barriers of crossing the monolayer.

### 3.1 Membrane-drug distance and number of contacts

Minimum distances between the different drug molecules and the mycolic acid monolayer have been presented in Fig. 2 for both the CHARMM and GROMOS force fields together with the snapshots of the systems at different time points (Snapshots of all the drug-monolayer systems at different time points have been presented in Fig. S1-S3 in SI). The minimum distance is defined as the minimum of all atomic-pair distances between the heavy atoms of drug molecules and those of the mycolic acid monolayer. For the combination of CHARMM force field parameters and ethambutol, the minimum distance first increased and, at ∼20ns, the distance decreased to 0.2 nm and remained stable for the rest of the 250 ns simulation time (Fig. 2(a)). This happened because of the preferential attachment of ethambutol to the tail of the mycolic acid chain. In case of GROMOS force field, the ethambutol molecule was attached to the monolayer surface within the first ∼300ps and remained there for the rest of the simulation window (Fig. 2(a)). Unlike CHARMM, ethambutol did not exhibit any preferential affinity toward the tail of the mycolic acid molecules, in the case of the GROMOS FF. More importantly, ethambutol never penetrated inside the mycolic acid monolayer but rather preferred to remain on the surface, for both CHARMM FF and GROMOS FF.

**Fig. 2:**
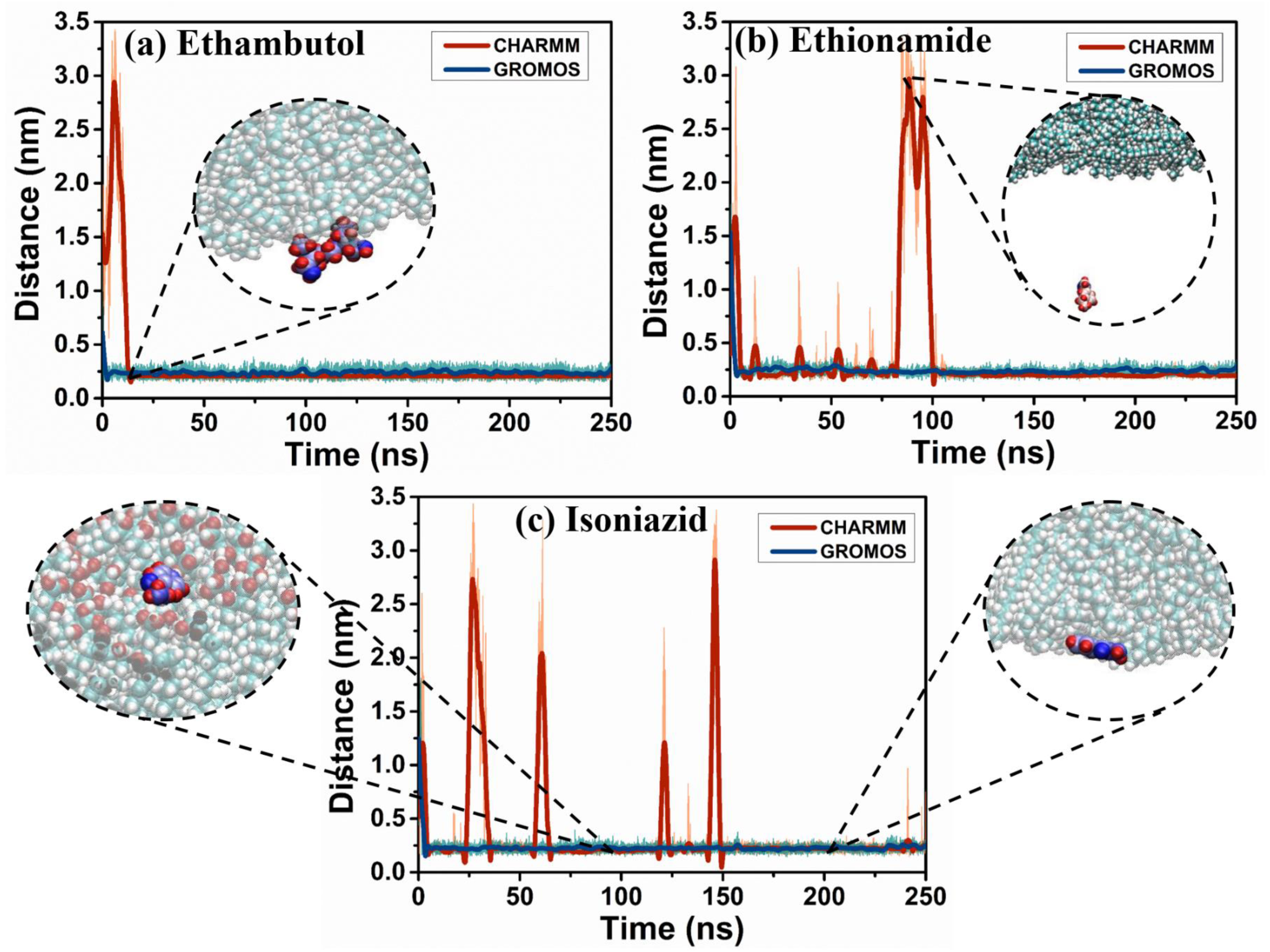
Minimum of the atomic pair distance or minimum distance is a signature of the drug-membrane interaction. Temporal evolution of the minimum distance between mycolic acid monolayer and (a) Ethambutol, (b) Ethionamide, and (c) Isoniazid for different FFs. Under GROMOS FF, the drug molecules are stably bound to the membrane surface. Snapshots of the drug-monolayer systems have been shown at different time point for the CHARMM FF with colour code for mycolic acid: C:Pale Blue, O: Red, H: white. Colour code for Drug molecule is based on atomic mass. For ethambutol/isoniazid: H:Red, O: Blue, N: light blue, C:whitish blue. For ethionamide: S:Deep Blue, H: red, C: reddish white, N: white.

A similar trend has been observed in the case of ethionamide. When CHARMM FF parameters were employed, the drug molecule attached to the mycolic acid monolayer within 5ns and subsequently periodically detached and again reattached to the membrane surface for the rest of the 250 ns long simulation (Fig. 2(b)). For the GROMOS FF, ethionamide was attached to the mycolic acid monolayer surface at ∼ 2ns and remained stable on the surface for the entire simulation window (Fig. 2(b)). For isoniazid as well, periodic adsorption and desorption of the drug molecule were recorded for CHARMM FF (Fig. 2(c)). When GROMOS FF parameters were used, isoniazid quickly got adsorbed stably on the surface of the monolayer (Fig. 2(c)). From the observed behavior, it can be commented that the GROMOS FF provides better stability of the drug molecules on the mycolic acid monolayer surface, compared to CHARMM FF.

We have also computed the number of contacts formed between the heavy atoms of the drug molecules and those in the mycolic acid monolayers. A contact is formed when any heavy atom of the drug molecule is within a distance of 0.4 nm of any heavy atom of mycolic acid. The number of contacts for various drug molecules and different force field parameters is presented in Fig.3. The number of contacts was generally noticed to be higher in the case of the GROMOS FF, for all of the drug molecules (Fig. 3). This reflects the fact that the drug molecules are more stable on the monolayer under GROMOS FF.

**Fig. 3:**
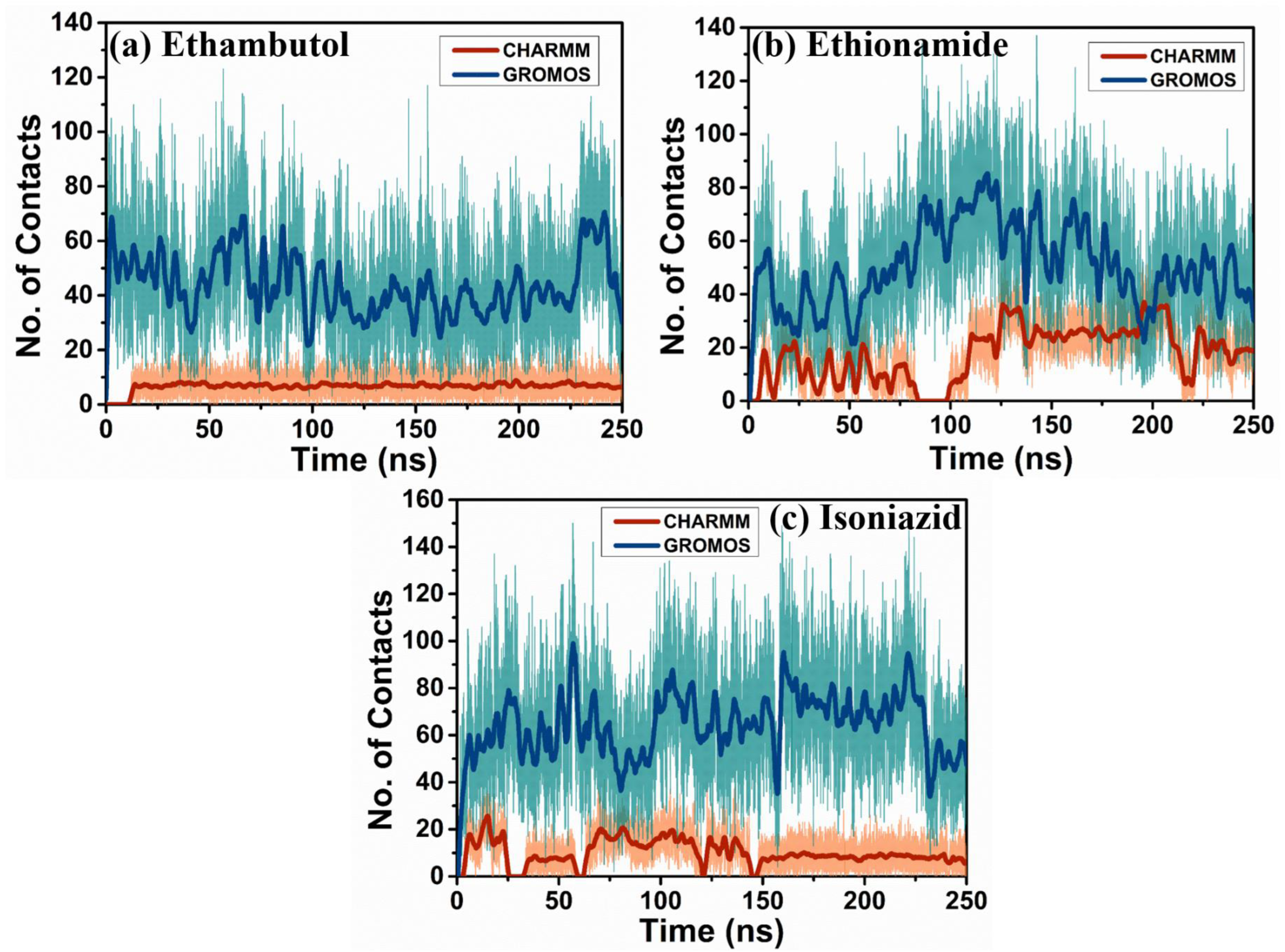
Number of contacts can be used to quantitatively assess the affinity of drug to membrane. Number of contacts formed between heavy atoms of mycolic acid monolayer and (a) Ethambutol, (b) Ethionamide, and (c) Isoniazid for different FFs. Number of contacts was found to be higher for GROMOS FF, confirming better stability of the drug molecules on monolayer surface.

### 3.2 Drug-monolayer interactions

The interactions of drug molecules with the mycolic acid monolayers is of utmost importance because it dictates the behavior of drug molecules in the membrane environment. In the present study, ethambutol was found to be interacting with the mycolic acid dominantly through VdW interactions, both for CHARMM and GROMOS FF (Fig. 4). Moreover, the VdW interaction energies were found to be of similar magnitudes for both set of force field parameters (Fig. 4). The coulombic contribution was noticed to be minor compared to VdW interactions, for both sets of force field parameters (Fig. 4).

**Fig. 4:**
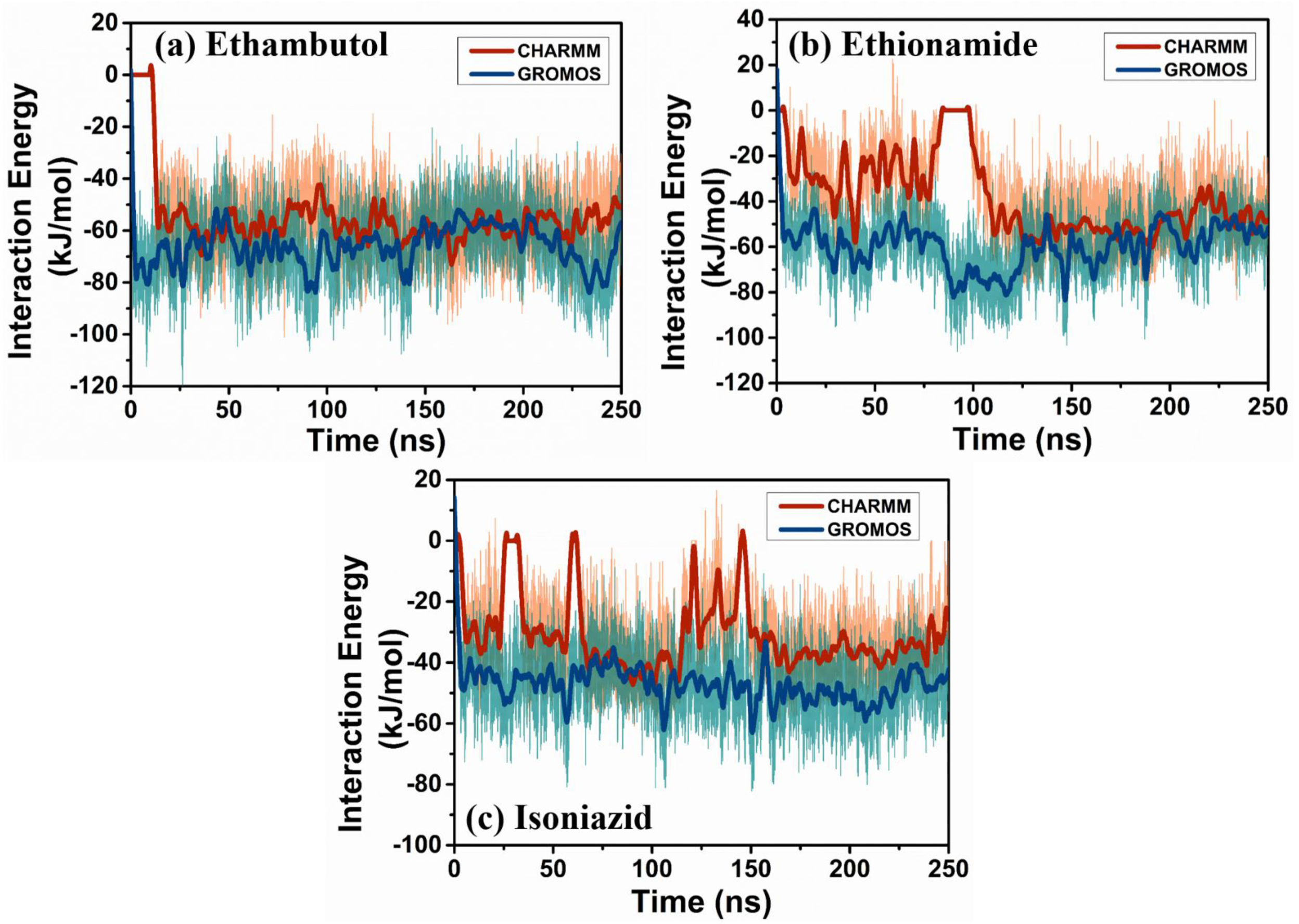
Different modes of interaction between drugs and monolayer dictate the system dynamics. Drug-monolayer VdW interaction energy as a function of time for (a) Ethambutol, (b) Ethionamide, and (c) Isoniazid. VdW interaction was found to be attractive in nature for all of the cases.

For ethionamide, when CHARMM FF was used, the drug molecule interacted with the monolayer via both electrostatic and VdW interactions, and the electrostatic interactions became dominant between 150ns and 200ns (Fig. 4(b) and 5(b)). For GROMOS FF, VdW interaction energy was noted to be the major contributor toward drug-membrane interactions, with negligible coulombic interactions (Fig. 4(b) and Fig. 5(b)). Although a weaker drug-membrane VdW interaction for CHARMM FF was observed for up to 125ns;, similar magnitudes were recorded afterward for both the FF (Fig. 4(b)). However, because of the difference in the magnitudes of the electrostatic interactions, the overall interactions between ethionamide and the mycolic acid membrane is significantly differs for CHARMM and GROMOS force field parameter sets.

**Fig. 5:**
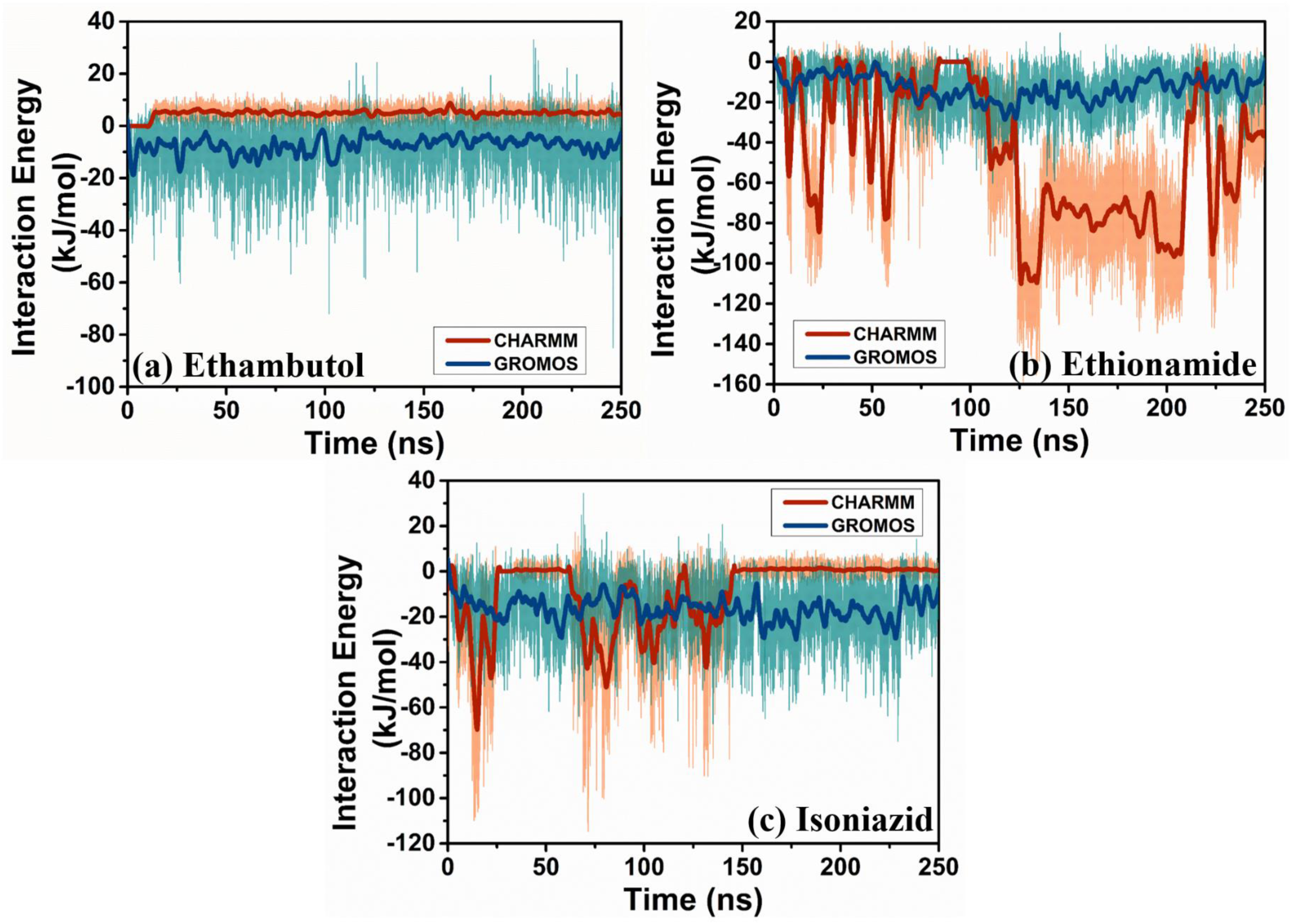
Coulombic interaction also influences behaviours of drug molecules. Drug-monolayer electrostatic interaction energy as a function of time for (a) Ethambutol, (b) Ethionamide, and (c) Isoniazid.

When isoniazid is considered, again the VdW interactions were found to be the dominant contributor for the drug-membrane interactions for both the CHARMM and GROMOS FF (Fig.4(c) and Fig. 5(c)). The VdW interactions was observed to be slightly greater for GROMOS FF throughout the simulation time (Fig. 4(c)), while the greater coulombic interaction was found for GROMOS force field from 150 ns onward (Fig.5(c)). From the above-mentioned behaviour, it can be easily concluded that GROMOS force field better stabilized the drug molecules on the mycolic acid monolayer membrane, compared to CHARMM FF. Our simulation study suggests that for both CHARMM and GROMOS FF, the strength of the drug-membrane VdW interactions for ethambutol was slightly greater than isoniazid (Fig. 4(a) and 4(c)), whereas the coulombic interaction for isoniazid was slightly higher for GROMOS FF (Fig. 5(a) and 5(c)).

Another important mode of interactions is the formation of hydrogen bonds between the drug molecules and the mycolic acids. Because of the fact that the oxygen atoms (receptors in the hydrogen bond formation) of ethambutol were exposed to the solvent side and there are only donor atoms present in the tail region of mycolic acids, this particular drug did not form any hydrogen bonds with the membrane, when CHARMM FF was used (Fig. 6(a)). The other two drug molecules (ethionamide and isoniazid) formed hydrogen bonds with the monolayer (Fig. 6(b) and 6(c)). It is noteworthy that isoniazid did not form H-bonds with monolayer after 150ns onward because after 150ns, isoniazid was attached to the tail side of the mycolic acid monolayer (see Fig. 2(c)), where donor/receptor atoms necessary for H-bond formation was absent (Fig. 6(c)). For GROMOS FF, all drug molecules formed hydrogen bonds with the monolayer and no significant differences was noted (Fig. 6).

**Fig. 6:**
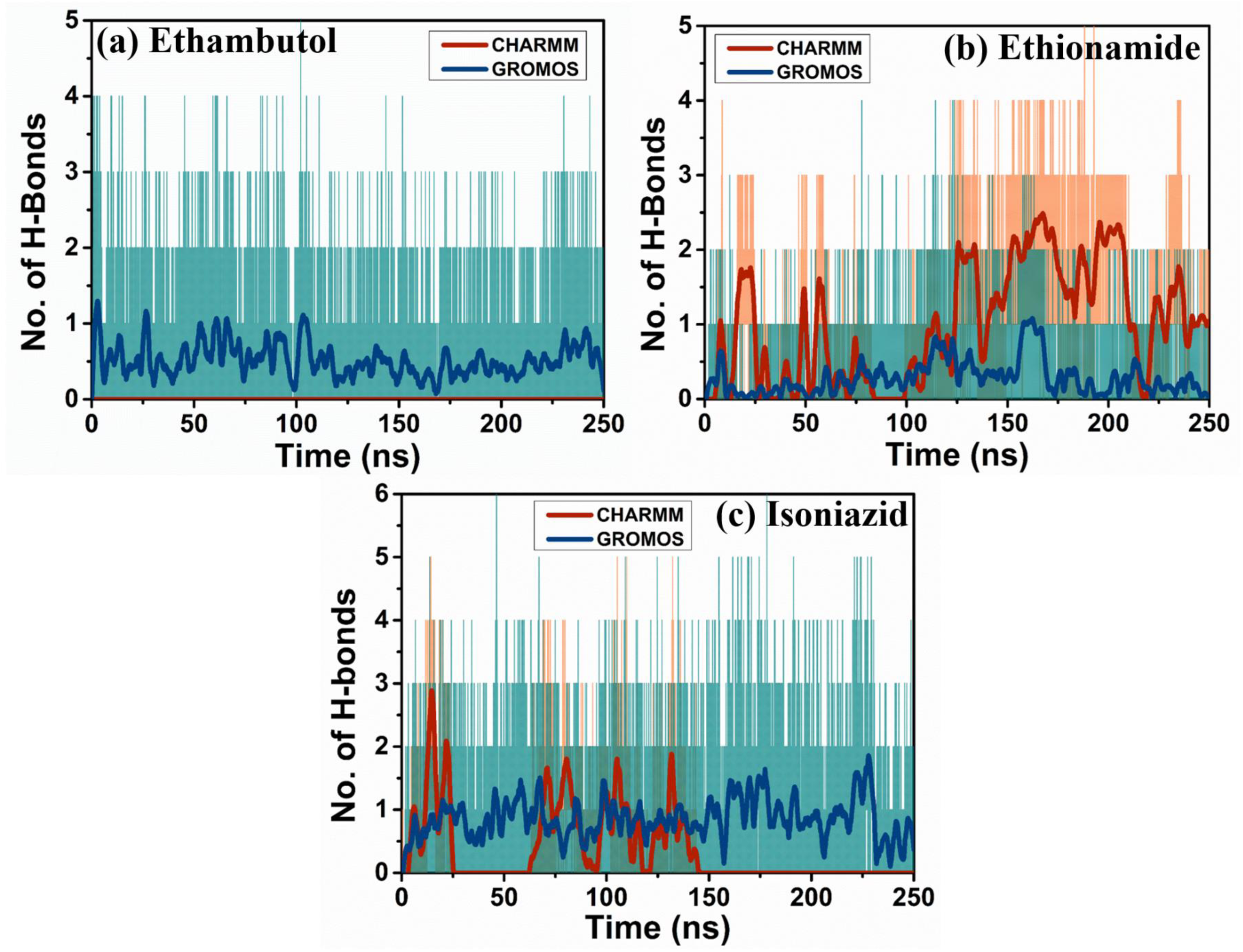
Hydrogen bond formed between drugs and monolayer surface affects the adsorption kinetics. Temporal behavior of the number of hydrogen bonds formed between the mycolic acid monolayer and (a) Ethambutol, (b) Ethionamide, and (c) Isoniazid, as a function of two force fields.

### 3.3 Drug-monolayer interactions during steered MD simulations

Different drugs interacted differently with the mycolic acid monolayer, while being pulled through the later. The various modes of the drug membrane interactions during pulling for two different force fields have been shown in Fig. 7. With the CHARMM force field, the primary mode of drug-membrane interaction is the VdW interactions (Fig. 7(a)). The strength of VdW interactions was found to be highest for ethambutol, followed by ethionamide, and isoniazid (Fig. 7(a)). For all the three drug molecules, the highest VdW interaction strength was recorded between ∼-1 nm and ∼2 nm distance from the COM of the monolayer, i.e around the middle portions of the mycolic acid assembly (Fig. 7(a)). The magnitude of VdW interaction energy decreased toward both ends of the mycolic acid layer (Fig. 7(a)).

**Fig. 7:**
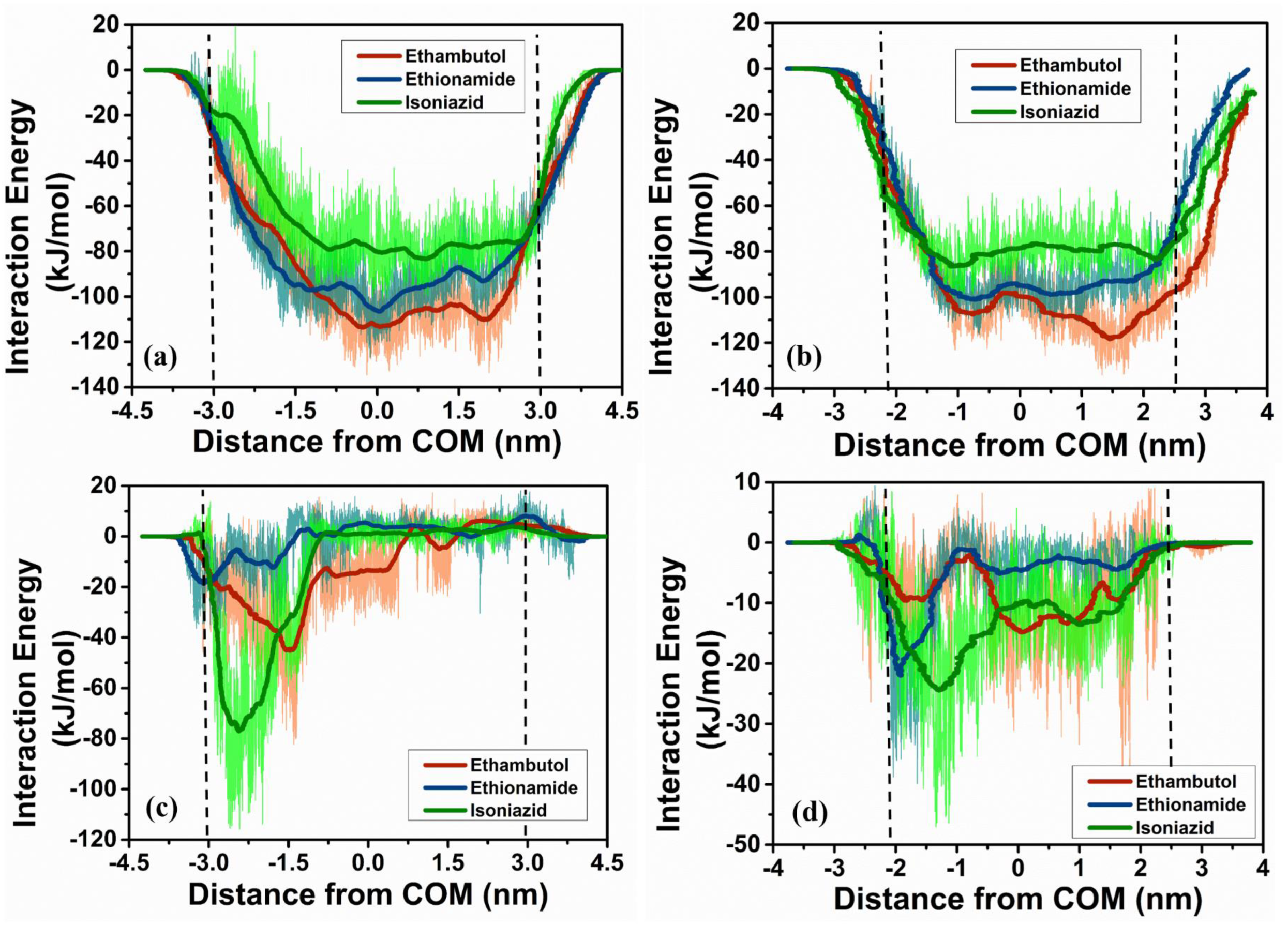
Different modes of drug-monolayer interaction during the pulling of drug molecules through the monolayer. Spatial variation of drug-membrane VdW interaction energy during the passage of drugs through the monolayer for (a) CHARMM and (b) GROMOS FFs. Positional dependency of electrostatic interactions for (c) CHARMM and (d) GROMOS FFs. In the figures, vertical black dashed lines represent positions of water-monolayer interfaces.

The nature of coulombic interactions were noticeably different for different drug molecules (Fig. 7(c)). Electrostatic interaction of isoniazid was highest in the head group region on the mycolic acid and became zero afterward (Fig. 7(c)). Ethionamide followed similar trend, but the strength of interaction was lesser (Fig. 7(c)). Ethambutol interacted with the membrane more strongly than ethionamide and unlike the other two molecules, it electrostatically interacted with the middle region of the membrane, as well (Fig. 7(c)). The hydrogen bond formation also followed the similar pattern. Isoniazid formed highest number of H-bonds with the head region of the monolayer but did not form any after that (Fig. S4(a) in SI). Ethambutol formed H-bonds with the head and middle portion of the monolayer while ethionamide formed H-bonds with every regions of the membrane (Fig. S4(a) in SI).

Similar trend in VdW interactions was observed with GROMOS parameters with the highest strength for ethambutol, followed by ethionamide and isoniazid (Fig. 7(b)). The coulombic interactions are also similar to CHARMM force fields, but for isoniazid, non-zero coulombic interaction is observed for every region of the monolayer (Fig. 7(d)). Moreover, Isoniazid formed hydrogen bonds with both head and tail region of mycolic acids with GROMOS force field (Fig. S4(b) in SI).

The strength of VdW interactions for each drug molecules were found to be very close for CHARMM and GROMOS parameters (Fig. 7(a)-7(b)). Significant differences have been observed in terms of electrostatic interactions. CHARMM offered a stronger coulombic interactions for ethambutol for head region of the monolayer, and slightly less electrostatic interactions for ethionamide (Fig. 7(c)). Most significant difference is noticed for isoniazid, which has been mentioned earlier. No significant change number of hydrogen bonds were probed for ethambutol and ethionamide with the change of the force field parameters (Fig. S4 in SI). Notable changes in the number of hydrogen bonds appeared for isoniazid, as mentioned earlier.

### 3.4 Free energy barrier and membrane permeability

The potential of the mean forces (PMF), obtained for all the three drugs from umbrella sampling using CHARMM FF, are plotted in Fig. 8(a). From the figure, it is evident that the free energy barrier is the highest for ethambutol (41.30 Kcal/mol), followed by ethionamide (39.39 Kcal/mol), and isoniazid (20.07 Kcal/mol). The free energy reached its minimum on the surface of the monolayer for all of the drug molecules, which, in corroboration with the other results, indicates that the drug molecules prefer to remain on the surface of the monolayer. The maxima of the free energy barrier for all three drugs were observed to be around the distance of 2 nm (Fig. 8(a)). It is worth stating that the order of the free energy barrier also followed the order of VdW interaction strength (Fig. 7(a) and 8(a)). Two peaks appeared in the free energy profile for every drug molecules, specifically for isoniazid because of the spatial change in the coulombic interactions and H-bond formation (Fig. 7(c) and S4(a) in SI), which are most prominently seen for isoniazid. Local minima can be seen ∼3nm distance (Fig. 8(a)), which is at the vicinity of the lower surface of the monolayer. These minima appeared because of the preference of the drug molecules to attach on the membrane surface.

**Fig. 8:**
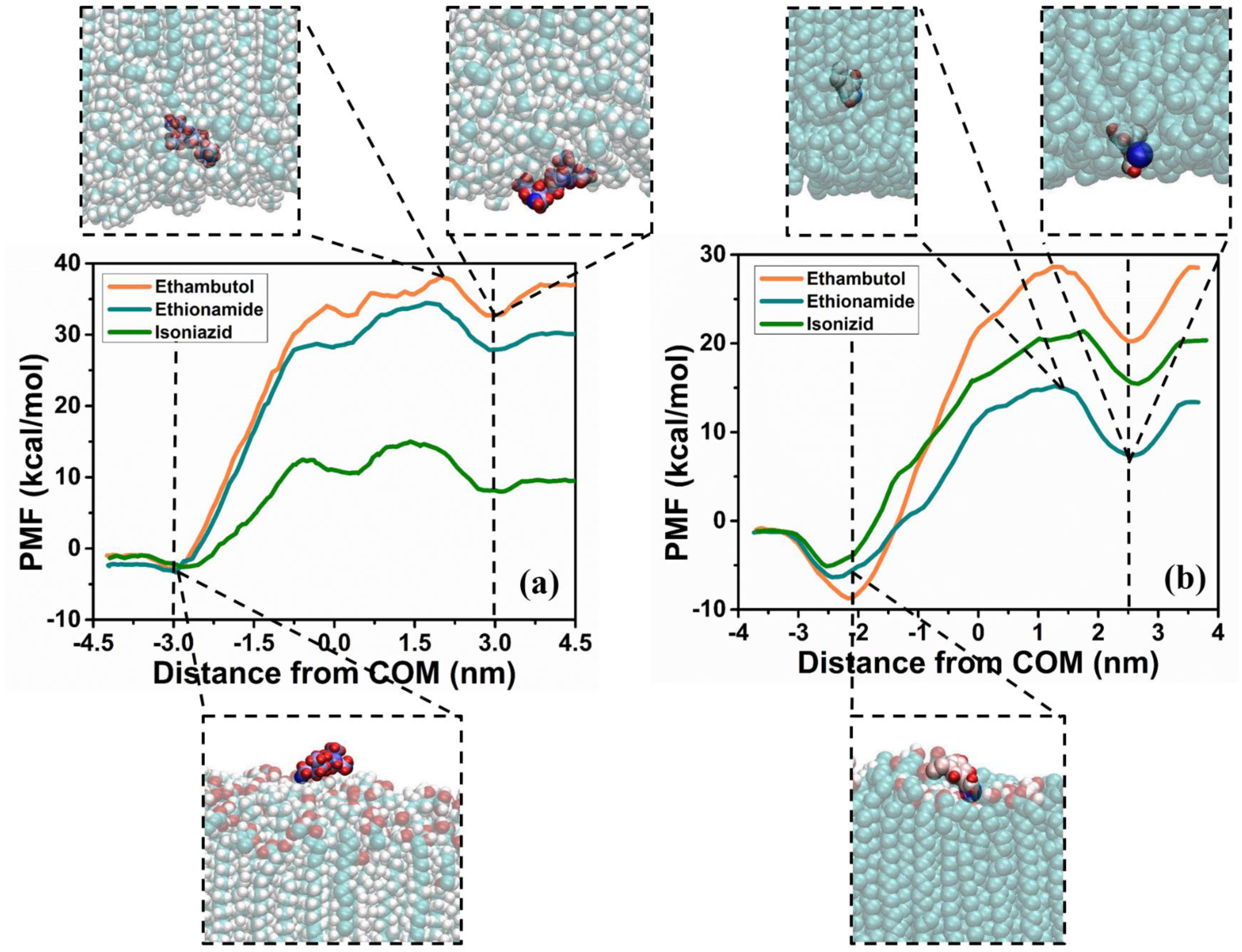
Mycolic acid monolayer offers high free energy barrier for drug molecules. Potential of mean force (PMF) for various pharmaceutical molecules for (a) CHARMM and (b) GROMOS FF. Free energy barrier was found to be higher for ethambutol and ethionamide for CHARMM FF, compared to GROMOS FF. Snapshots of different drug-monolayer systems have depicted at various points of PMF curve, with colour code for mycolic acid: C:Pale Blue, O: Red, H: white. Colour code for Drug molecule is based on atomic mass. For ethambutol: H:Red, O: Blue, N: light blue, C:whitish blue. For ethionamide: S:Deep Blue, H: red, C: reddish white, N: white. In the figures, vertical black dashed lines represent positions of water-monolayer interfaces.

The shape of the PMF profiles of the different drug molecules for GROMOS parameters have been plotted in Fig. 8(b). The major differences of GROMOS PMF profiles with the CHRAMM PMF profiles are the following.

1. With GROMOS parameters, although the ethambutol exhibited the highest free energy barrier (37.79 Kcal/mol) like CHARMM; it is followed by isoniazid (27.99 Kcal/mol), and ethionamide (21.78 Kcal/mol) (fig.9©).
2. Two distinct peaks appeared in the CHARMM PMF profile is not noticed for GROMOS parameters.

**Fig. 9:**
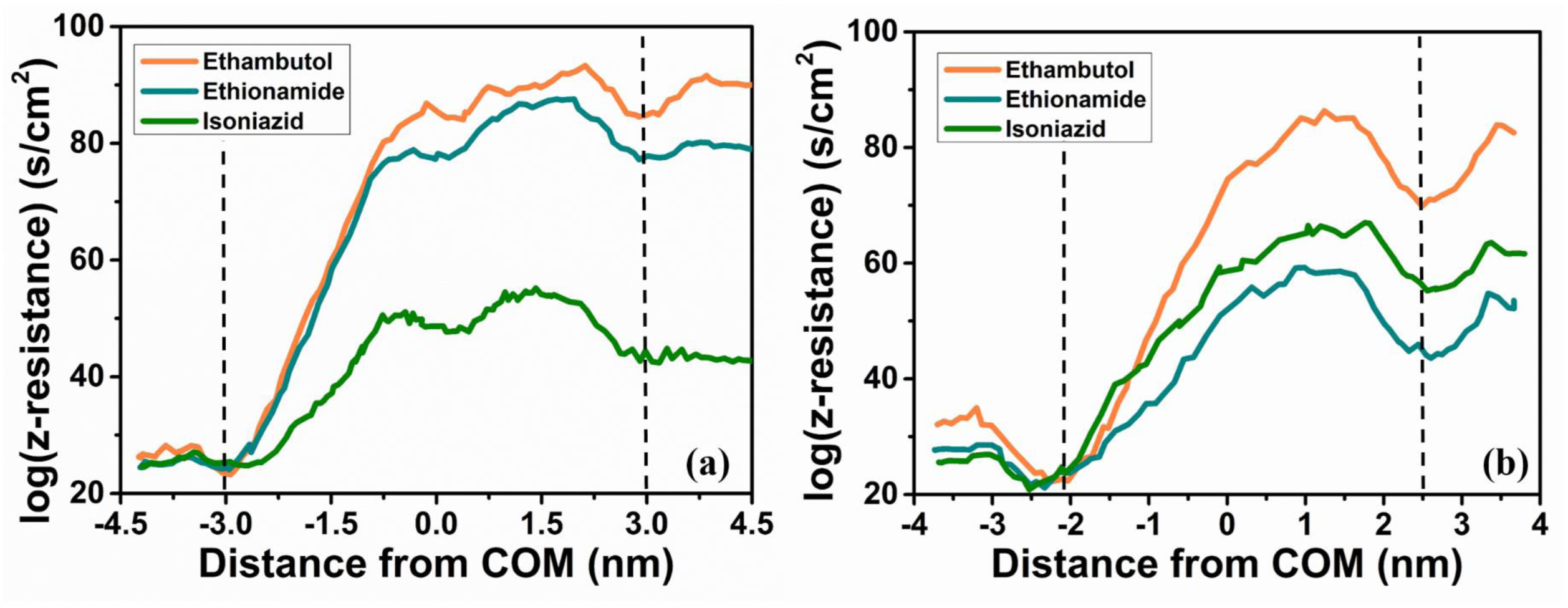
Mycolic acid membrane is responsible for low drug penetration property of TB bacterium. Resistance of mycolic acid monolayer against different drug molecules for (a) CHARMM and (b) GROMOS FF. Higher resistance recorded for ethambutol and ethionamide for CHARMM FF, compared to GROMOS FF. In the figures, vertical black dashed lines represent positions of water-monolayer interfaces.

The other characteristics of the GROMOS PMF profiles are similar to that of CHARMM. For instance, the affinity of drug molecules to the membrane surfaces can be clearly seen from GROMOS PMF profiles. Besides this, the height of the free energy generally followed the spatial strength of the drug-membrane VdW interactions (Fig. 7(b)). However, although the strength of isoniazid-mycolic acid VdW interactions was slightly lower than that between ethionamide and membrane, the free energy barrier for the former is higher (Fig. 8(b)).

The disappearance of the dual peaks in the GROMACS PMF profiles can be attributed to the significantly altered spatial profile of coulombic interactions and number of H-bonds, compared to CHARMM parameters (Fig. 7(c)-7(d) and Fig. S4 in SI). More importantly, from Fig. 7(d), it is clear that the electrostatic interactions between isoniazid and mycolic acid is stronger than that between ethionamide and mycolic acid monolayer in the tail region of the molecules. This significantly affected the PMF profile and lower the free energy barrier of ethionamide, than isoniazid. This feature is absent for CHARMM parameters and thus, the free energy barrier for ethionamide was higher than isoniazid.

The position-dependent resistance of the mycolic acid membrane against all the three drug molecules, which follows the same pattern as the free energy barrier, has been represented using Log scale in Fig. 9, which followed the similar patter like the PMF profile (Fig. 8), for both the FFs. The diffusion constant has been presented in Fig. 10, and it shows that the diffusivity is extremely low for the mycolic acid monolayer, in agreement with the existing literature.^34^ The effective permeability of the membrane for different drug molecules has been tabulated in Table 1 for both CHARMM and GROMOS FF. For CHARMM force field parameters, isoniazid was found to be most effective in penetrating the mycolic acid monolayer, and ethambutol is the least effective. However, ethionamide was found to be more effective in terms of permeability under GROMOS FF (Table 1). It should be noted that the effective permeability was found to be higher for ethambutol and ethionamide for GROMOS FF (Table 1). Besides that, the position dependent diffusion constant was always calculated to be higher for GROMOS FF for all of the drug molecules (Fig. S5 in SI), which is also a reason for the higher permeability of ethambutol with GROMOS parameters (Table 1).

**Fig. 10:**
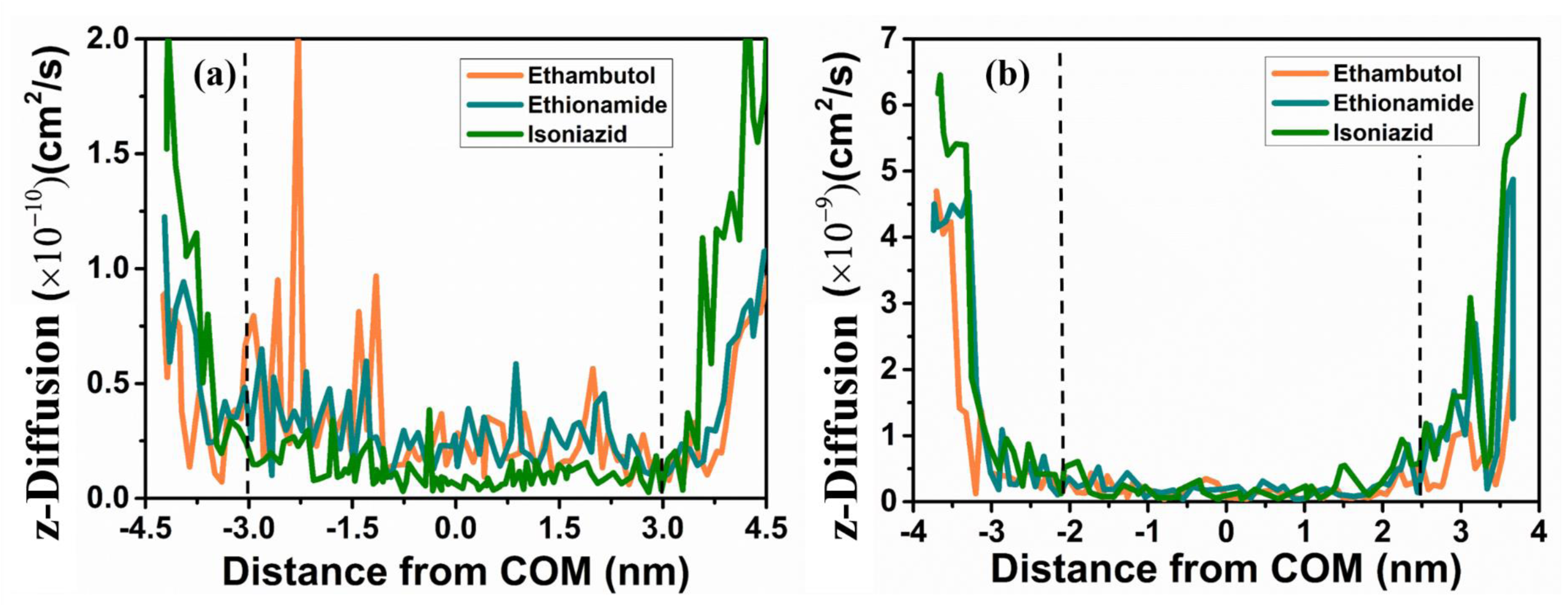
Mycolic acid monolayer acts as a diffusion barrier for drug molecules. Position-dependent diffusion coefficients of various pharmaceutical molecules in the mycolic acid monolayer for (a) CHARMM and (b) GROMOS FFs. In the figures, vertical black dashed lines represent positions of water-monolayer interfaces.

**Table 1:** Effective permeability (cm/s) of different drug molecules using two different FF: CHARMM36 all atom FF (CHARMM) and GROMOS 54A7-ATB (GROMOS) united-atom FF.

| Name of the drugs | GROMOS | CHARMM |
| --- | --- | --- |
| Ethambutol | $9.61 \times 10^{-31}$ | $8.40 \times 10^{-34}$ |
| Ethionamide | $3.57 \times 10^{-19}$ | $1.45 \times 10^{-31}$ |
| Isoniazid | $1.68 \times 10^{-22}$ | $2.94 \times 10^{-17}$ |

## 4. Discussion

In the present current study, we have observed that the drug molecules differed in terms of the affinity toward the monolayer surface as a function of FF parameters. In the case of ethionamide and isoniazid, GROMOS FF provided better stability of the drug molecules on the monolayer surface, whereas the affinity of ethambutol toward the monolayer surface was found to be similar for both the FF (Fig. 2). The underlying reasons can be explained in the following manner.

From Fig. 4, it is clear that the VdW interaction was similar for both sets of FFs in the case of ethambutol almost, but weaker for CHARMM FF for ethionamide for the first ∼125ns. For isoniazid, VdW interaction was found to be weaker for CHARMM FF for almost throughout the simulation window (Fig. 4(c)). This weaker VdW interaction may contribute to the less stable dynamics, observed for ethionamide and isoniazid under CHARMM FF. One interesting observation is that ethionamide electrostatically interacted with the monolayer (with CHARMM FF), while the magnitude of coulombic interaction for the other two (ethambutol and isoniazid) is not that significant (Fig. 5). This strong electrostatic interaction helped ethionamide to have more stability on the monolayer surface, compared to isoniazid under CHARMM FF, as seen from the evolution of the minimum distance (Fig. 2(b)-2(c)).

While comparing the dynamics of ethionamide using two different FFs, one interesting aspect that appears is that despite having greater coulombic interaction, Ethionamide was found to have less affinity toward the monolayer under CHARMM FF. To figure out the underlying reason, we have computed the radial distribution function (g(r)) of water oxygen atoms surrounding the drug molecules to assess the hydration effect on the dynamics of the drug molecules, and the calculated g(r) for both sets of FFs have been presented in Fig. S5 of SI. From Fig. S5, it can be clearly seen that under CHARMM FF, ethionamide and isoniazid had been surrounded by greater number of water molecules, compared to GROMOS FF. Higher solvation contributed to the less stable behaviour of ethionamide and isoniazid on mycolic acid monolayer surface under CHARMM FF, despite having greater drug-monolayer electrostatic interaction (for ethionamide). The solvation behaviour was quite similar for ethambutol for both sets of FFs (Fig. S5(a) in SI), and it exhibited similar affinity toward the monolayer surface for CHARMM and GROMOS.

It is a well-known fact that GROMOS FF, in many instances, overestimated surface area/lipid for various kinds of lipids like DOPC, DLPC, DMPC and POPC, and such overestimation led to enhanced water penetration in the membrane.^35^ In the present study, area per lipid heads group was found to be higher for GROMOS FF (Fig. S6 in SI). This gives rise to higher diffusivity for ethambutol, when GROMOS FF is used. For the other two molecules, several other factor like the drug-monolayer interactions played an important role in determining the diffusivity, and that reflected in the observed trend in permeability, as presented in Table 1.

Apart from the above-mentioned facts, we have computed the electrostatic potential and electric field across the membrane for both the FF in absence of any drug molecule, in order to obtain a better insight into the behavior of the drug molecules inside the membrane, and they have been presented in Fig.S7 of the SI. The electrostatic potential was found to be higher at the water-monolayer interface(Fig. S7(a) in SI), which, we believe, helped to better stabilize the drug molecules on the surface, as well (Fig. 2). Also, for GROMOS FF, electrostatic potential changed more smoothly inside the membrane, compared to that for CHARMM FF. This difference also may have played a role behind the notable difference in PMF profile of the drug molecules, discussed earlier. Other important fact is the drastic difference in the effective membrane permeability for different drug molecules, which arise from the exponential dependency of effective permeability on the free energy profile (see section 2), because of which slight change in PMF profile results in drastic change in effective permeability.

It is a well-known fact that the transport across membrane takes place through two major ways: active (proteins are involved), and passive (diffusion of small molecules across the membrane).^36^ Relatively hydrophobic molecules can passively transport across the membrane.^36^ In the current study, the possibility of passive transport of some first line tuberculosis drugs has been explored using two different well know FF. Among the three tested drugs, ethambutol exhibited free highest energy barrier while using both the FF. For the other two molecules, further in-depth study or experimental evidences is required to reach a conclusion about the accuracy of the force field parameters. In summary, we can say that, in terms of drug-monolayer interactions, both the force field provided consistent results, significant differences were observed in the thermodynamic aspects of the drug transport and the adsorption of the drugs on the monolayer surface.

## 5. Conclusions

In the current study, we have successfully employed atomistic MD simulation to investigate drug-mycolic acid monolayer interactions using two different FF: namely GROMOS and CHARMM. We have compared the biding affinity of different drug molecules with the mycolic acid monolayer and have estimated the drug permeability through the monolayer from the free energy profile. Our The following key conclusions can be drawn from the present work.

a) Drug-monolayer interaction energies were found to be comparable for both the CHARMM and GROMOS FF. For both the FFs, drug molecules interacted majorly via VdW interactions.
b) For both the FFs, the drug molecules preferred to stay on the monolayer surface. However, GROMOS FF provided better stability of the drug molecules, specifically ethionamide and isoniazid because of the lesser solvation and higher electrostatic potential at the water-membrane interface. On the other hand, ethionamide exhibited affinity toward the tail of the mycolic acid monolayer for adsorption, which was not observed for simulations using GROMOS FF.
c) The PMF profile of different drug molecules looked significantly different for GROMOS and CHARMM FF because of the notable difference of drug-mycolic acid coulombic interaction within the monolayer (during passage of the drugs through the monolayer). The free energy barrier height also differed significantly under the two FFs because of the same reason. However, ethambutol was found to experience the highest free energy barrier for monolayer crossing for both the FFs.
d) The height of free energy barrier was recorded to be lower for GROMOS FF, compared to that for CHARMM FF because of the higher area per head group of lipid for GROMOS FF.
e) Our simulation results indicate that the drugs will typically take time in the order of milliseconds to cross the monolayer through passive diffusion.

Summarizing, the current work explored the interactions of some of the first line tuberculosis drugs with mycolic acid monolayer and establishes the importance of the choice of the force field parameter sets in carrying out such study. More importantly, we believe that the current study paves a pathway for future *in silico* studies on the interplay between potential TB drug candidates and M.Tb cell wall.

## Supporting information

file contains additional figures

## Acknowledgements

The authors thank the Department of Biotechnology (DBT), Government of India, for funding this work (Grant number: BT/PR33123/MED/29/1497/2020). The authors also thank Prof. Narayanaswamy Jayaraman, Prof. Dipankar Chatterji, and Dr. Anirban Ghosh, IISc, Bangalore, for critical reading of the manuscript and useful suggestions.

