## Supplementary material for "Drug-bacterial membrane interactions: A Tale of two force fields": file contains additional figures

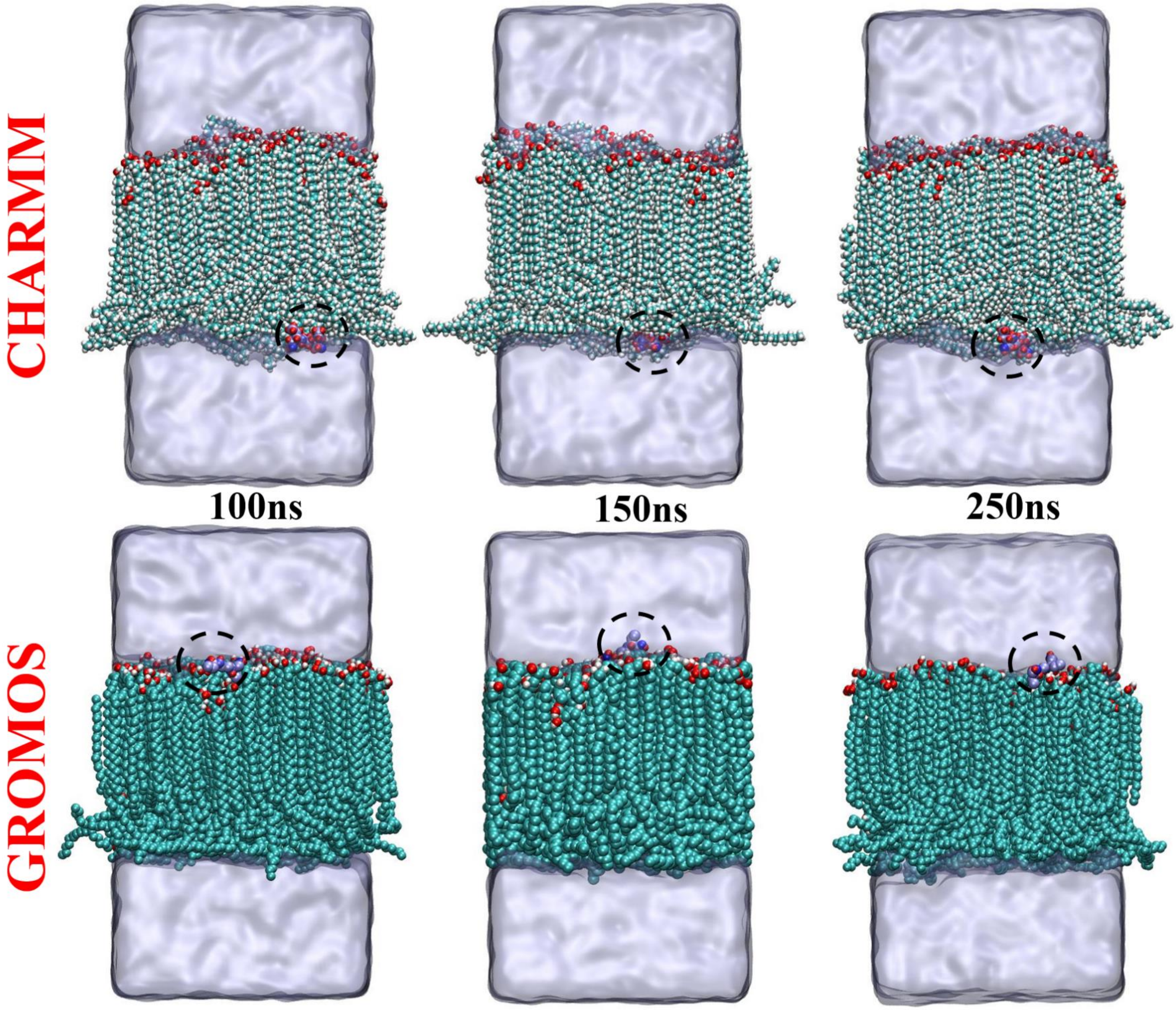


**Fig. S1:** snapshots of ethambutol-mycolic acid monolayer system at different time points. Colour code for mycolic acid: C:Pale Blue, O: Red, H: white. Colour code for Drug molecule is based on atomic mass. For ethambutol: H:Red, O: Blue, N: light blue, C:whitish blue. Water is presented using ice-blue transparent surface. Drug molecule is encircled using dashed line for clarity.

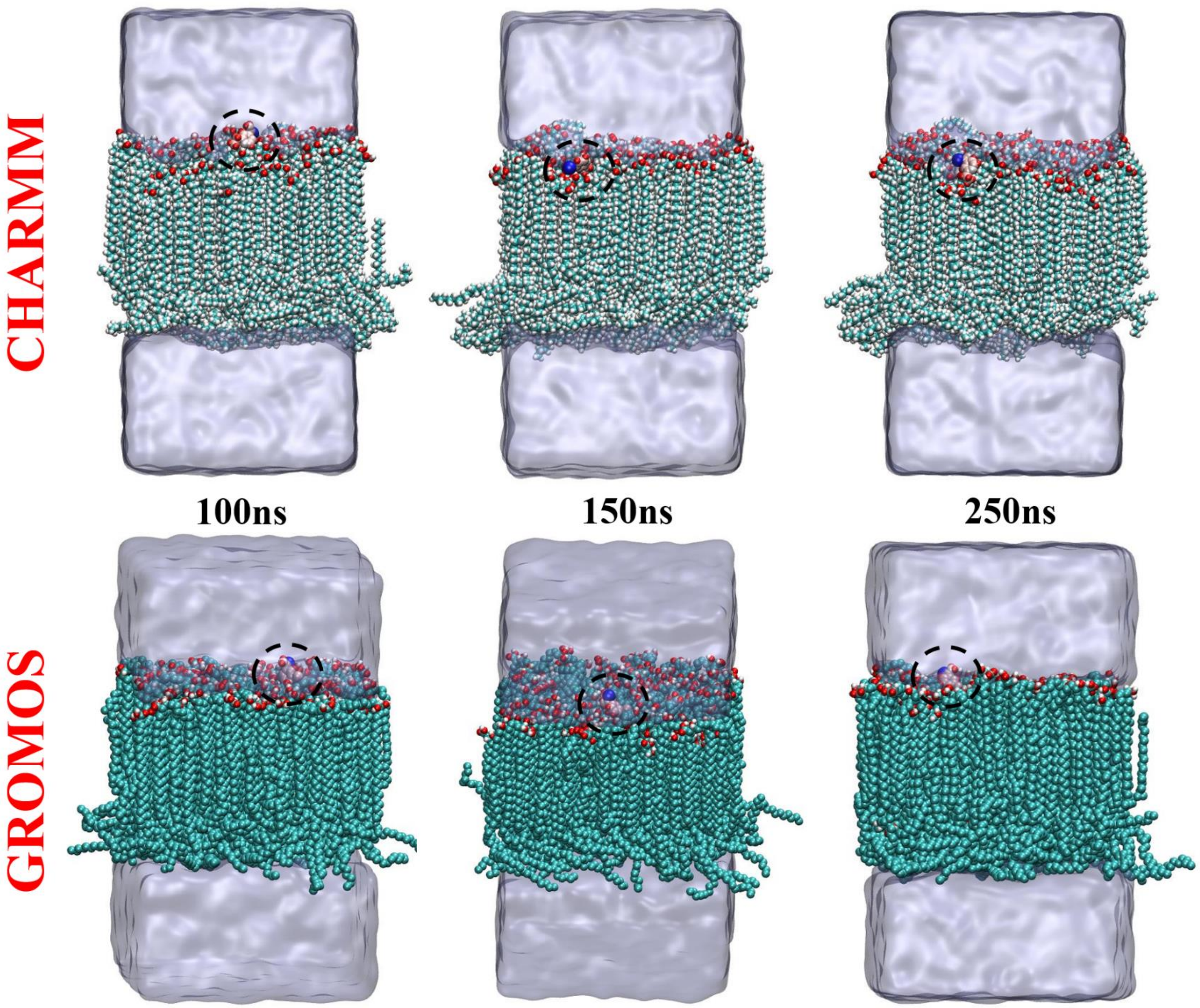


**Fig. S2:** snapshots of ethionamide-mycolic acid monolayer system at different time points. Colour code for mycolic acid: C:Pale Blue, O: Red, H: white. Colour code for Drug molecule is based on atomic mass. For ethionamide: S:Deep Blue, H: red, C: reddish white, N: white. Water is presented using ice-blue transparent surface. Drug molecule is encircled using dashed line for clarity.

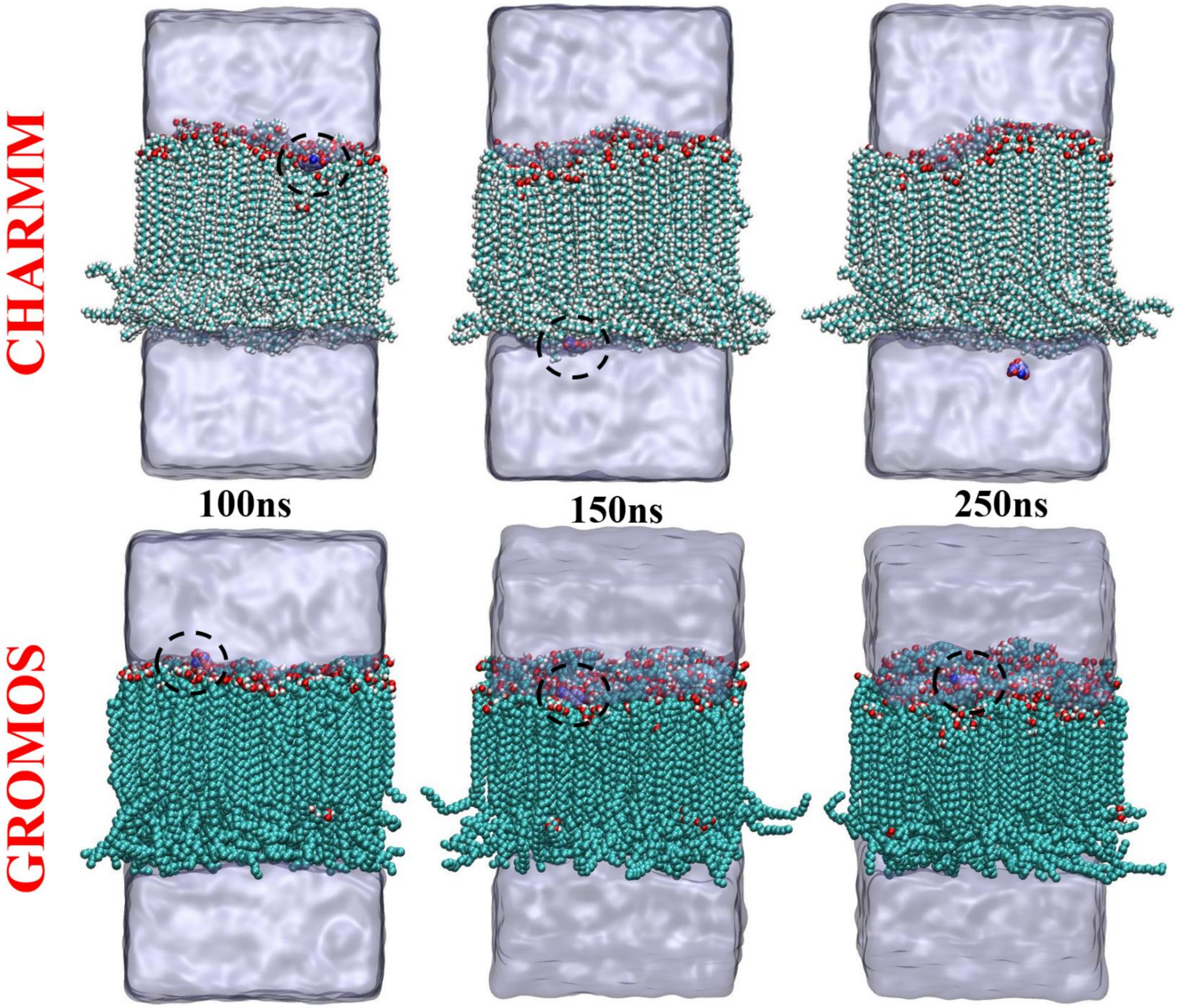


**Fig. S3:** snapshots of Isoniazid-mycolic acid monolayer system at different time points. Colour code for mycolic acid: C:Pale Blue, O: Red, H: white. Colour code for Drug molecule is based on atomic mass. For isoniazid: H:Red, O: Blue, N: light blue, C:whitish blue. Water is presented using ice-blue transparent surface. Drug molecule is encircled using dashed line for clarity.

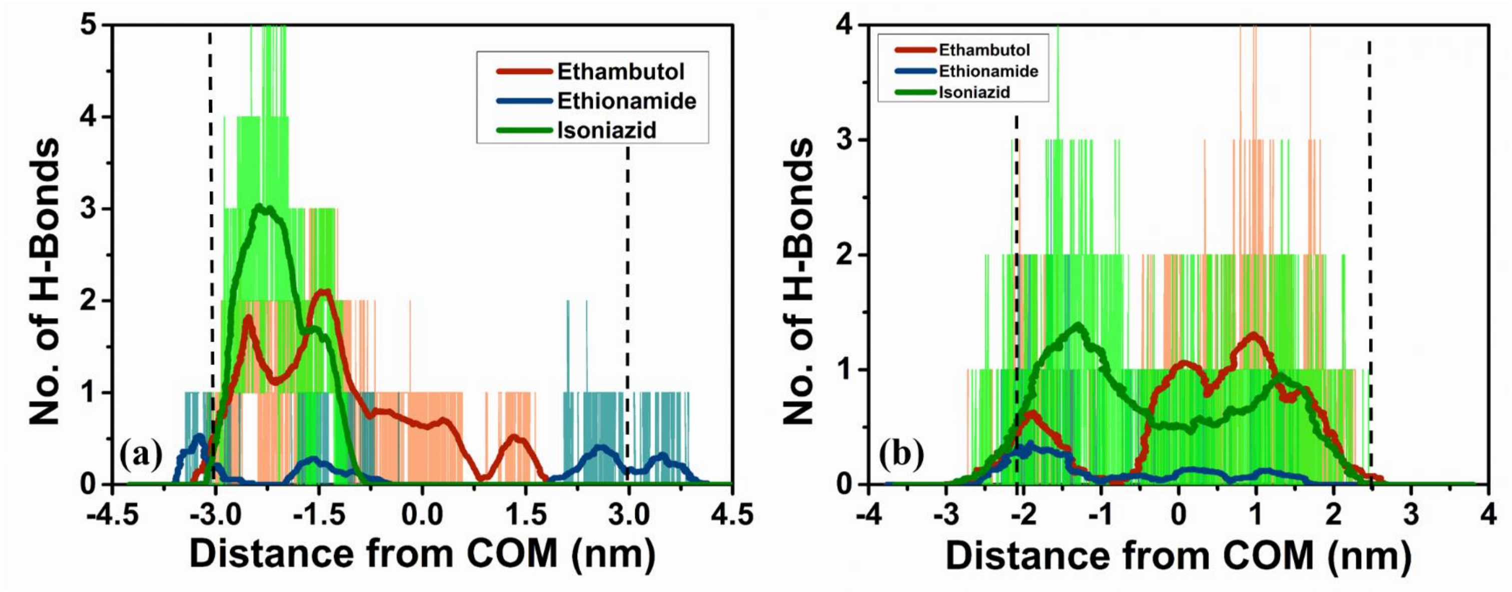


**Fig. S4:** Spatial variance of the number of hydrogen bonds during the pulling of drug molecules through the monolayer for (a) CHARMM and (b) GROMOS FFs.

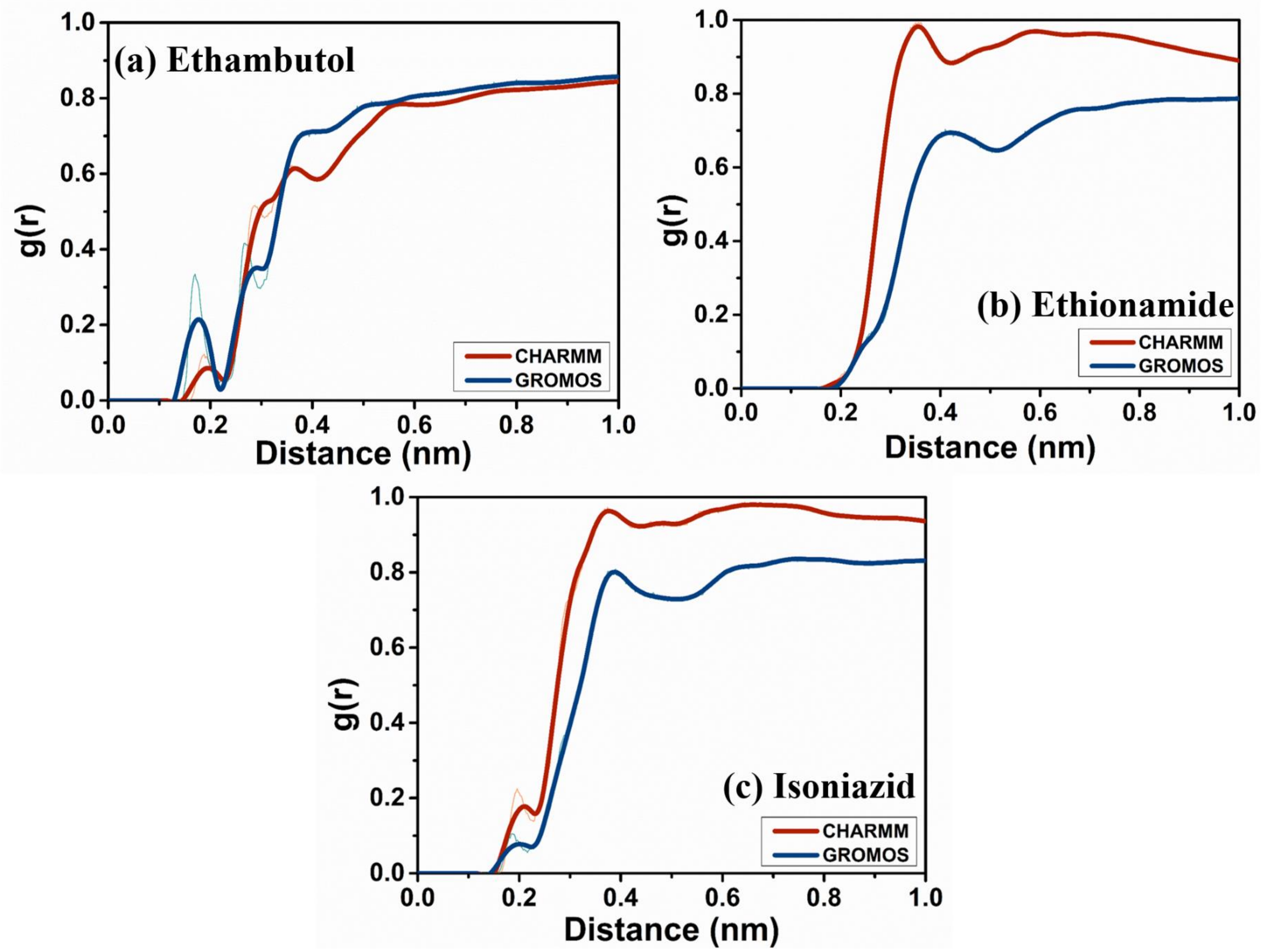


**Fig. S5:** Radial distribution function of water oxygen atoms w.r.t. centre of mass of (a) Ethambutol, (b) Ethionamide, and (c) Isoniazid for different FFs.

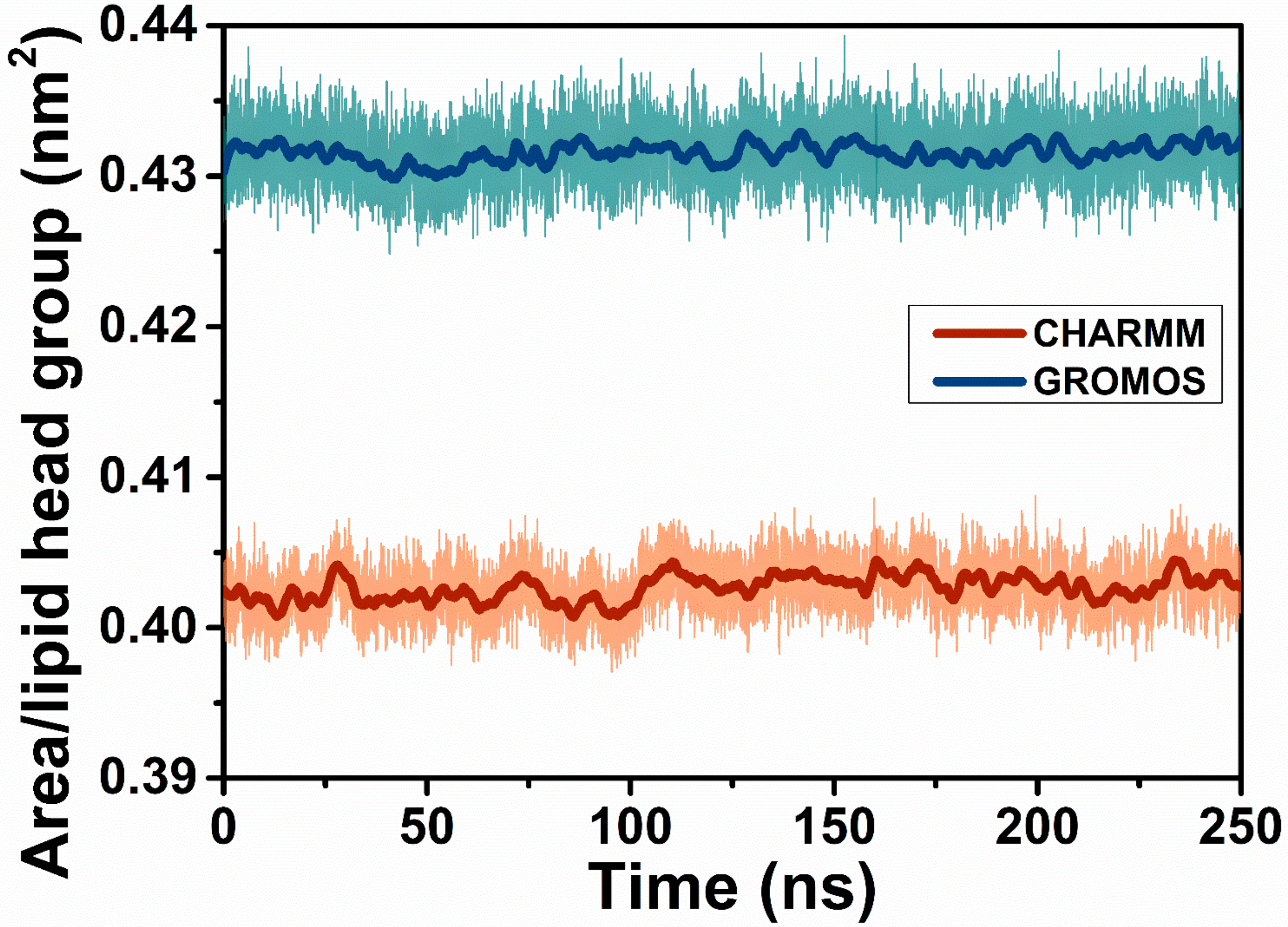


**Fig. S6:** Area/lipid of the mycolic acid monolayer for two different FFs.

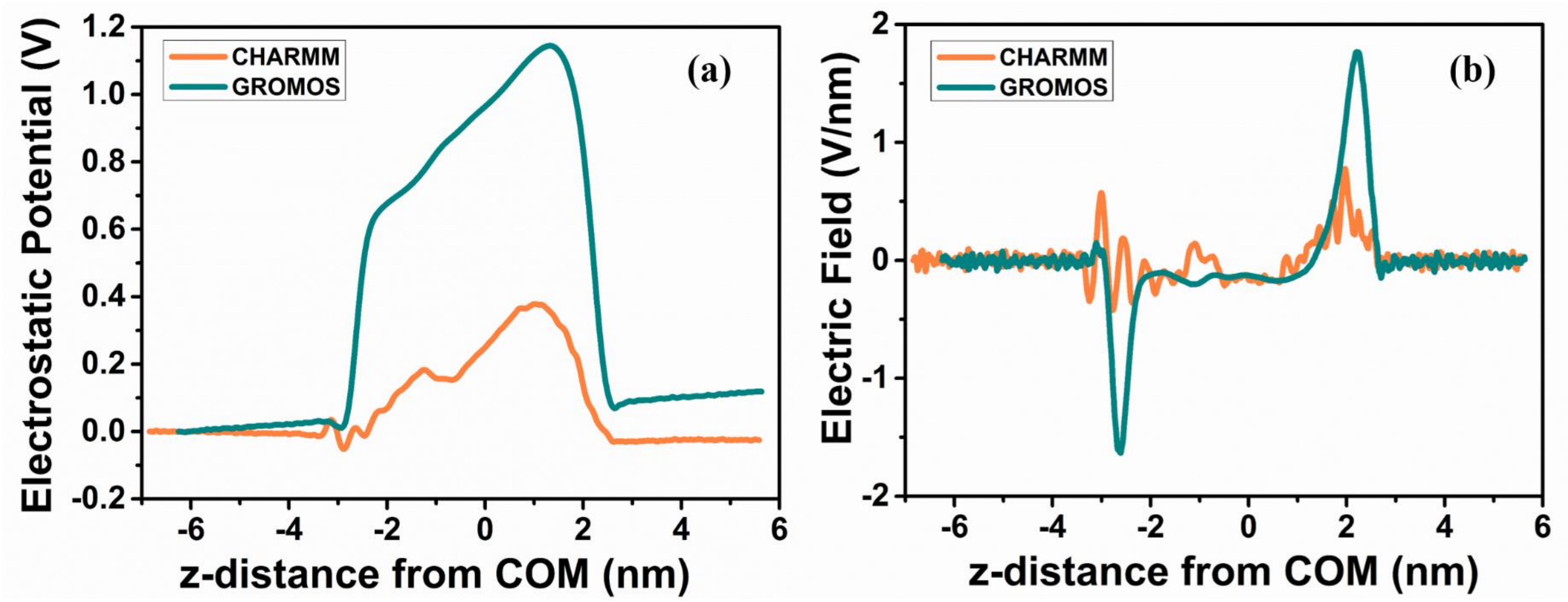


**Fig. S7:** (a) Electrostatic potential and (b) electric field intensity across the z-direction of the simulation box for two different FFs.
